# The +1 nucleosome functions in Pol II transcription initiation and the transition to elongation

**DOI:** 10.64898/2026.05.11.724227

**Authors:** Yumeng Zhan, Julio Abril-Garrido, Frauke Grabbe, Paulina Seweryn, Ute Neef, Christian Dienemann, Patrick Cramer

**Author notes:** These authors contributed equally: Yumeng Zhan, Julio Abril-Garrido.

## Abstract

Transcription initiation by RNA polymerase II (Pol II) occurs next to a +1 nucleosome that is positioned downstream of the transcription start site (TSS). The +1 nucleosome has been shown to influence the pre-initiation complex (PIC) assembly and Pol II pausing, but it is unclear whether and how it functions in transcription initiation and the transition to elongation. Here, we investigate the transcription initiation-elongation transition *in vitro* using DNA templates containing a +1 nucleosome, and we present cryo-EM structures of five intermediate states. First, after PIC assembly, the +1 nucleosome evicts TFIID from the PIC upon binding of ATP to TFIIH. Second, after DNA opening, the +1 nucleosome stimulates TFIIH translocase activity and initial RNA synthesis. Finally, after DNA bubble rewinding, the +1 nucleosome removes TFIIH from the early elongation complex for promoter escape. Our findings show that the +1 nucleosome not only acts as a passive border for PIC assembly and a passive barrier for Pol II pausing, but rather has active functions during the initiation-elongation transition of transcription.

## INTRODUCTION

RNA polymerase II (Pol II) transcribes all protein-coding and some non-coding genes. Before transcription initiates, the coactivators Mediator and TFIID and a set of general transcription factors (GTFs: TFIIA, TFIIB, TFIIE, TFIIF and TFIIH) recruit Pol II to the gene promoter, which together with Pol II form the Pol II transcription pre-initiation complex (PIC)^1–4^. Sequence elements^5^ such as the TATA-box, initiator (Inr), motif ten element (MTE) and downstream promoter elements (DPE) guide PIC assembly at the promoter, and the combination of these elements influences the efficiency of the assembly^6^.

After PIC assembly, the TFIIH subunit XPB hydrolyses ATP and translocates on the downstream DNA to generate torque that leads to DNA duplex melting^7,8^, establishing the open complex (OC)^7,8^. Transcription initiation by Pol II (synthesis of the first phosphodiester bond) converts the OC to an initially transcribing complex (ITC). Due to its unstable nature, the ITC is prone to abortive transcription and therefore relies on TFIIH and further ATP hydrolysis for efficient RNA synthesis^9–11^. As Pol II elongates the RNA, the ITC becomes more stable^9,10^ but generates an overextended initial transcription bubble^9,11^ by DNA scrunching^12–14^. When the RNA reaches a length of 10-14 nt, the upstream end of the overextended bubble rewinds, resulting in a smaller but more stable transcription bubble that converts the ITC to an elongation complex (EC)^9,11,15–17^. It has been shown that transcription bubble rewinding coincides with the eviction of TFIIB, TFIIA and TBP from the transcription complex^16,17^. TFIID, however, has been suggested to remain bound to the transcription complex after bubble rewinding, and its dissociation might depend on the downstream DNA sequence^17^. TFIIE and TFIIH can stay associated with the early EC and can be displaced upon recruitment of the elongation and pausing factors DSIF and NELF^16^, which completes promoter escape by transcribing Pol II^15,16,18^. At this point, the transcribing complex completes the transition to elongation, a process encompassing the synthesis of the second phosphodiester bond to promoter escape.

Gene promoters generally contain a well-positioned nucleosome immediately downstream of the transcription start site (TSS) that is commonly known as the +1 nucleosome^19–23^. The +1 nucleosome has been shown to influence PIC assembly and TSS selection. In yeast, histone-DNA interactions can bury the TSS ∼10 bp inside the +1 nucleosome^24,25^, which impairs TFIIH translocase activity during TSS scanning^13,26–29^ and thereby alters TSS selection^30^. At mammalian promoters, the position of the +1 nucleosome differs depending on gene expression levels. For inactive genes, the edge of the +1 nucleosome is positioned at ∼10 bp downstream of the TSS^19,31,32^, which prevents TFIIH from engaging with promoter DNA and keeps TFIIH in a closed inactive conformation^31^. Promoters of active genes, however, possess a +1 nucleosome that is located at ∼40-50 bp downstream of the TSS^19,24,31–33^. Recent studies have shown that a +1 nucleosome positioned 40-50 bp downstream of the TSS can interact with Mediator and TFIIH^34^ and have suggested it may facilitate PIC assembly on genes without core promoter elements such as the TATA-box^3,6,17,35–37^.

The intimate association of the +1 nucleosome with the PIC during PIC assembly suggests that the +1 nucleosome may also influence transcription initiation and the transition to elongation, but there is currently no evidence for this. The mechanism of PIC assembly in the context of a +1 nucleosome has been studied structurally and biochemically^30,31,34^. However, insights into the mechanism of transcription initiation and the transition to elongation are limited to studies performed on linear promoter DNA^7–18,38–42^, leaving the role of the +1 nucleosome in these processes poorly understood.

In this work, we combine biochemical assays with structural studies to investigate a possible functional role of the +1 nucleosome in the transcription initiation-elongation transition. We demonstrate that the +1 nucleosome stimulates initial transcription by regulating TFIIH activity and that it controls the timely release of TFIID and TFIIH from the transcription complex. Together with cryo-EM structures of five intermediates during transcription initiation and the transition to elongation in the presence of the +1 nucleosome, our study establishes the functional importance of the +1 nucleosome for transcription initiation and the transition to elongation and its molecular basis.

## RESULTS

### Binding of ADP-BeF_3_ to XPB destabilizes TFIID from the PIC in the presence of a +1 nucleosome

Immediately following the assembly of the PIC, the promoter DNA is melted by the ATP-dependent translocase activity of TFIIH subunit XPB. To investigate the role of the +1 nucleosome in this process, we designed a promoter DNA template that positions the edge of the nucleosome 40 bp downstream of the TSS (Figure 1A, Methods), a distance consistent with active gene promoters^19,24,31–33^. After assembling the nucleosome on the promoter DNA, we formed a PIC using highly purified human GTFs: TFIIA, -IIB, -IID, -IIE, - IIF and -IIH and *Sus scrofa* Pol II, which is 99.9% identical to human RNA Pol II (Figure S1A). Finally, we performed cryo-EM analysis on this PIC-nucleosome complex in the absence (PIC-Nuc^40W^) or in the presence of the ATP analogue ADP-BeF_3_ that locks the XPB ATPase in the pre-hydrolysis state (PIC-Nuc^40W^_ADP-BeF3_).

**Figure 1.**
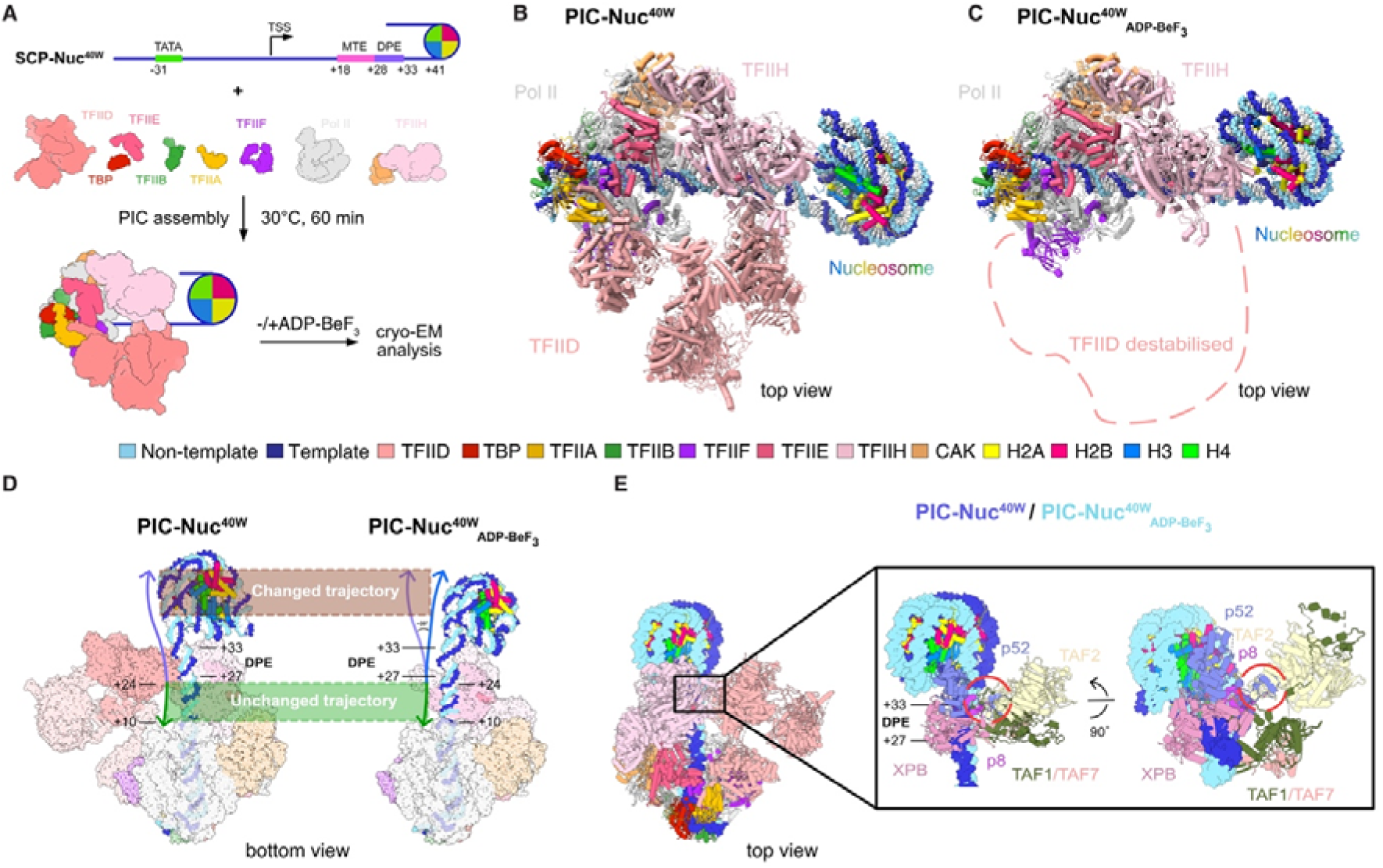
Binding of ADP-BeF_3_ to XPB destabilizes TFIID from the PIC in the presence of a +1 nucleosome. **(A)** Scheme of the experimental setup for cryo-EM sample preparation of the PIC-Nuc^40W^ and the PIC-Nuc^40W^_ADP-BeF_ structures shown in **(B)** and **(C)**, using the SCP-Nuc^40W^ DNA template. Positions of core promoter motifs and the edge of the +1 nucleosome are indicated relative to the TSS. **(B)** Structure of the PIC-Nuc^40W^ complex without ADP-BeF_3_, shown as cartoon in top view. DNA is represented as spheres. **(C)** Structure of the PIC-Nuc^40W^_ADP-BeF_ shown as cartoon in top view. DNA is represented as spheres. Dashed line indicates where TFIID was in the PIC-Nuc^40W^. **(D)** Side-by side comparison of the PIC-Nuc^40W^ and the PIC-Nuc^40W^_ADP-BeF_ in bottom view as atom-sphere representation. Purple and blue arrows highlight the trajectory of the PIC-Nuc^40W^ and the PIC-Nuc^40W^_ADP-BeF_, respectively. Regions in the promoter with unchanged (+10 to +24) and changed trajectory (DPE - +27 onwards), and their specific position with respect to the TSS, are depicted with dashed lines. **(E)** Overlay of the PIC-Nuc^40W^ and the PIC-Nuc^40W^_ADP-BeF_ structures in top view aligned on the DPE motif. The TAF2 APD and TFIIH p52-p8 dimerization domain clash because of the changed promoter DNA trajectory. The clash is highlighted with a red circle. Parts of TFIID and TFIIH were removed for clarity.

We obtained a reconstruction for the PIC-Nuc^40W^ at an overall resolution of 8.5 Å and focused classifications and refinements resolved the subregions of core PIC (cPIC), TFIIH, TFIID and the nucleosome to 4.4 Å, 11.1 Å, 12.3 Å and 7.5 Å, respectively (Methods, Figure S1B and 2A). Cryo-EM analysis of the PIC-Nuc^40W^_ADP-BeF3_ led to a reconstruction at an overall resolution of 4.2 Å, with local resolutions of 3.6 Å, 4.2 Å, 3.1 Å and 4.1 Å for cPIC, TFIIH, XPB and nucleosome, respectively (Methods, Figure S1C and 2B). We found clear density for ADP-BeF_3_ in the nucleotide binding site of XPB in the PIC-Nuc^40W^_ADP-BeF3_ (Figure S2C), confirming that this structure represents the nucleotide bound state of XPB.

In the absence of ADP-BeF_3_ (PIC-Nuc^40W^), the reconstruction of PIC-Nuc^40W^ showed density for all GTFs and the overall PIC conformation resembles previously published PIC structures with nearby +1 nucleosomes^34,43^ (Figure 1B and Figure S2D). In contrast, when ADP-BeF_3_ was present (PIC-Nuc^40W^_ADP-BeF3_) we did not observe density for TFIID in the reconstruction (Figure 1C, Figure S2E). This suggests that TFIID is destabilized upon the binding of ADP-BeF_3_ to TFIIH. Previous studies have shown that TFIID remains bound to promoter DNA in an ITC without a +1 nucleosome^17^, indicating that the destabilization of TFIID by ADP-BeF_3_ binding to XPB requires the +1 nucleosome. ATPases such as XPB have been shown to translocate on the DNA approximately 1 bp upon binding to ATP^38,44,45^. Indeed, in PIC-Nuc^40W^_ADP-BeF3_ we found the +1 nucleosome rotated towards TFIIH and the MTE-DPE motif shifted away from TBP-associated factors (TAFs) 1 and 7 (TAF1 and TAF7) (Figure 1D). This sterically prevents the binding of TFIID to the MTE-DPE motif as this would lead to a clash between p52/p8 of TFIIH and TAF2 (Figure 1E). The rotation of the +1 nucleosome also establishes a contact between TFIIH subunit p52 and the nucleosome, which has been described before^31,34^ (Figure S2F). This likely reduces flexibility of the downstream DNA and thereby prevents the MTE-DPE motif from moving back towards TAF1/7.

Altogether, these structures show that binding of the ATP analogue ADP-BeF_3_ to TFIIH subunit XPB changes the path of the promoter DNA and the orientation of the +1 nucleosome relative to the PIC, which displaces TFIID from the MTE-DPE motif. Additionally, positioning of TFIIH subunits p52/p8 in the PIC-Nuc^40W^_ADP-BeF_ prevents TFIID from rebinding. Thus, in the context of a +1 nucleosome, binding of ADP-BeF_3_ to XPB leads to destabilization of TFIID and may eventually evict it from the PIC.

### The +1 nucleosome contacts TFIIH during initial transcription

To investigate how initial RNA synthesis functions in the presence of the +1 nucleosome, we proceeded to structurally characterize the initially transcribing intermediates obtained by *de novo* transcription on promoter DNA templates containing a +1 nucleosome. We modified the promoter DNA and introduced U2 or U6-less cassettes to stall Pol II transcription after synthesis of 2 or 6 nt, respectively (Figure 2A, Methods). After reconstituting the +1 nucleosome on this promoter DNA, we assembled the PIC and initiated transcription with adenosine triphosphate (ATP) and cytidine triphosphate (CTP) for the U2-less cassette and ATP, CTP and guanosine triphosphate (GTP) for the U6-less cassette, and analysed the resulting transcription reactions using cryo-EM (Figure 2A and Methods). We obtained cryo-EM reconstructions of the resulting complexes at overall resolutions of 7.5 Å and 7.6 Å, respectively (Figure 2B-C, Figure S3A-B and Figure S4A-B). We name the two structures ITC2-Nuc^40W^ and ITC6-Nuc^40W^ since focused refinements of the Pol II active site showed the presence of 2 and 6 nt RNAs, respectively (Figure 2B-C, inserts).

**Figure 2.**
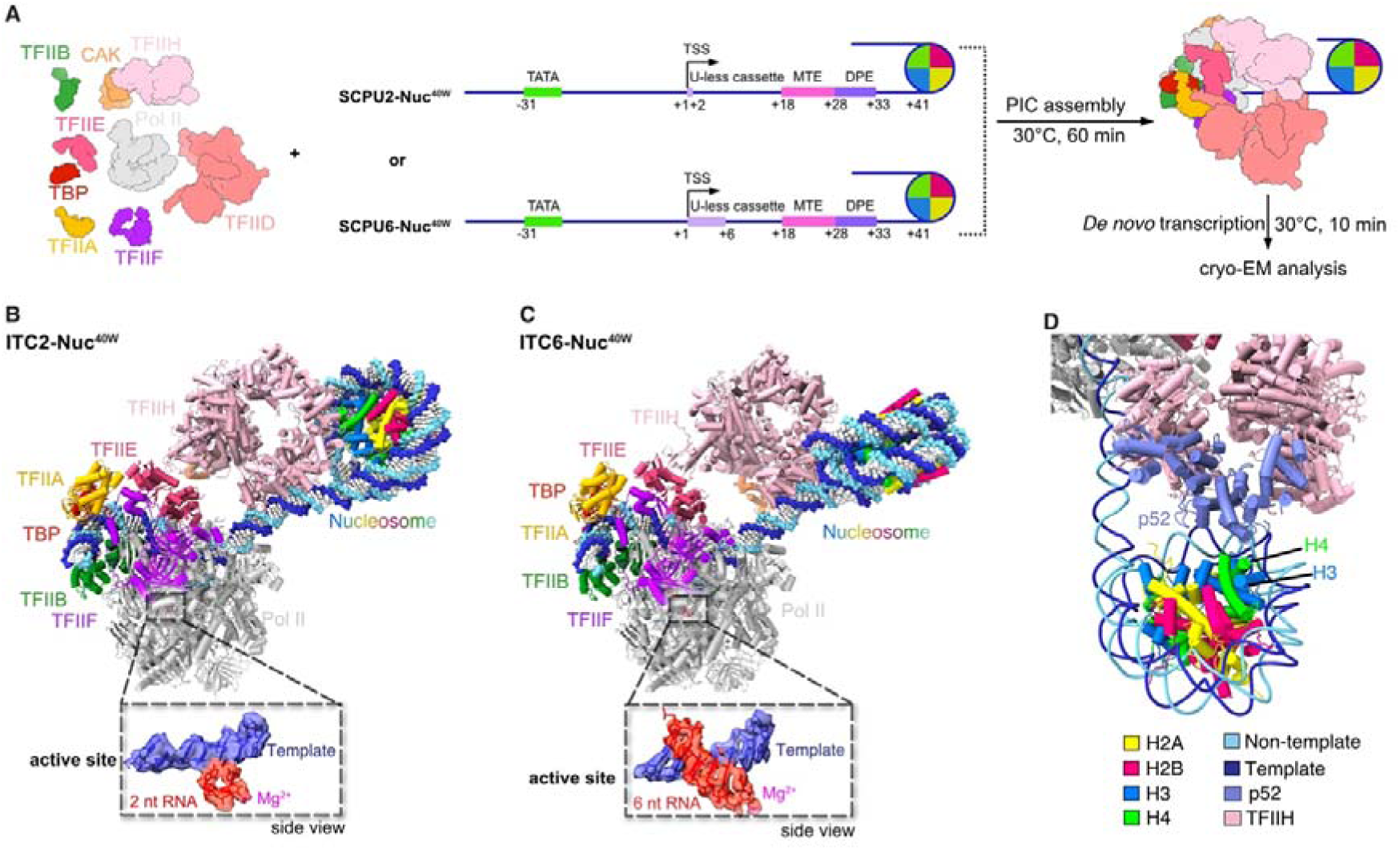
TFIIH contacts the +1 nucleosome during initial transcription. **(A)** Scheme of the experimental setup for cryo-EM sample preparation of the ITC2-Nuc^40W^ and the ITC6-Nuc^40W^ structures shown in (**B)** and (**C)**, using the SCPU2-Nuc^40W^ and the SCPU6-Nuc^40W^ DNA templates, respectively. Positions of core promoter motifs, U-less cassette and the edge of the +1 nucleosome are indicated relative to the TSS. **(B),** Structure of the ITC2-Nuc^40W^ shown as cartoon in side view, the DNA is shown as spheres. The inset below depicts the cryo-EM density carved around the Pol II active site showing that Pol II has transcribed 2 nt RNA in the ITC2-Nuc^40W^. The active site Mg^2+^ is shown as a magenta sphere. **(C),** Structure of the ITC6-Nuc^40W^ shown as cartoon in side view, the DNA is shown as spheres. The inset below depicts the cryo-EM density carved around the Pol II active site showing that Pol II has transcribed 6 nt RNA in the ITC6-Nuc^40W^. The active site Mg^2+^ is shown as a magenta sphere. **(D)** The TFIIH-nucleosome interface in the ITC2-Nuc^40W^. The +1 nucleosome contacts TFIIH subunit p52 via H3 and H4.

Both structures contain density for TBP, TFIIA, -IIB, -IIE, -IIF and -IIH (Figure S4C). We did not observe density for TFIID in the ITC2-Nuc^40W^ or the ITC6-Nuc^40W^, consistent with the proposed TFIID destabilization and eviction upon binding to the ATP analogue ADP-BeF_3_. Compared to the PIC-Nuc^40W^_ADP-BeF_, TFIIH and the +1 nucleosome shift and rotate relative to cPIC in the ITC2-Nuc^40W^, likely to accommodate transcription of 2 nt (Figure S4D), whilst the p52-nucleosome interaction is maintained (Figure 2D). When transcription proceeded in the ITC6-Nuc^40W^, the nucleosome rotates by ∼150° compared to the ITC2-Nuc^40W^ (Figure S4E), which is consistent with the rotational advancement expected from the DNA helical geometry that transcribing an additional 4 nt would introduce. Despite the large rotation of the nucleosome, the nucleosomal DNA still interacts with the region between p52 and XPB (Figure S4F), similar to what has been observed for the PIC before^34^. 3D multibody refinement shows that both the nucleosome and TFIIH can rotate and shift relative to the cPIC. However, despite their flexibility relative to the cPIC, TFIIH and the nucleosome always contact each other. (Figure S4G).

Altogether, our cryo-EM structures show that the +1 nucleosome establishes interactions with TFIIH upon binding to ATP and the nucleosome remains bound to TFIIH during formation of the first phosphodiester bond. When transcription proceeds to 6 nt, the nucleosome rotates but still contacts TFIIH subunits XPB and p52. Thus, the +1 nucleosome contacts TFIIH during initial transcription, at least up to 6 nt of RNA.

### The +1 nucleosome stimulates TFIIH activity and initial RNA synthesis

The TFIIH subunit XPB is essential for translocating promoter DNA during promoter DNA melting and initial transcription^7,8,46,47^ and p52 was shown to regulate ATPase activity of XPB^48,49^. Therefore, our observation that the +1 nucleosome interacts with XPB and p52 during initial transcription prompted us to test whether a +1 nucleosome positioned 40 bp downstream of the TSS has an influence on TFIIH activity in the context of a PIC.

To this end, we first performed an NADH-coupled ATPase assay to monitor dATP hydrolysis by TFIIH, in the context of a PIC assembled on either linear promoter DNA or promoter DNA with a +1 nucleosome positioned 40 bp downstream of the TSS (Figure 3A). Indeed, the ATPase activity of TFIIH is ∼1.5 fold higher when a +1 nucleosome is present on promoter DNA (Figure 3A). Next, we asked if stimulation of the ATPase activity by the +1 nucleosome also results in enhanced DNA translocase activity. For that, a 22-bp triplex was positioned immediately downstream of where XPB binds promoter DNA in the closed PIC (Figure 3B), and triplex displacement was visualized by gel electrophoresis. Upon addition of ATP, CTP and GTP, we observe increased triplex displacement when a +1 nucleosome is present compared to linear DNA (Figure 3C), which is consistent with stimulated ATPase activity in the context of a +1 nucleosome. Because the translocase activity of TFIIH is crucial for initial transcription^7,8,46,47^, we also tested if the +1 nucleosome impacts initial transcription (Figure 3D). Indeed, a +1 nucleosome positioned at 40 bp downstream of the TSS stimulates production of short RNAs of 6 and 7 nt from a U-less DNA template (Figure 3E). This stimulation is absent when the nucleosome is positioned 70 bp downstream of the TSS (Figure 3E), suggesting that the +1 nucleosome has to be spatially close to the PIC to stimulate initial transcription.

**Figure 3.**
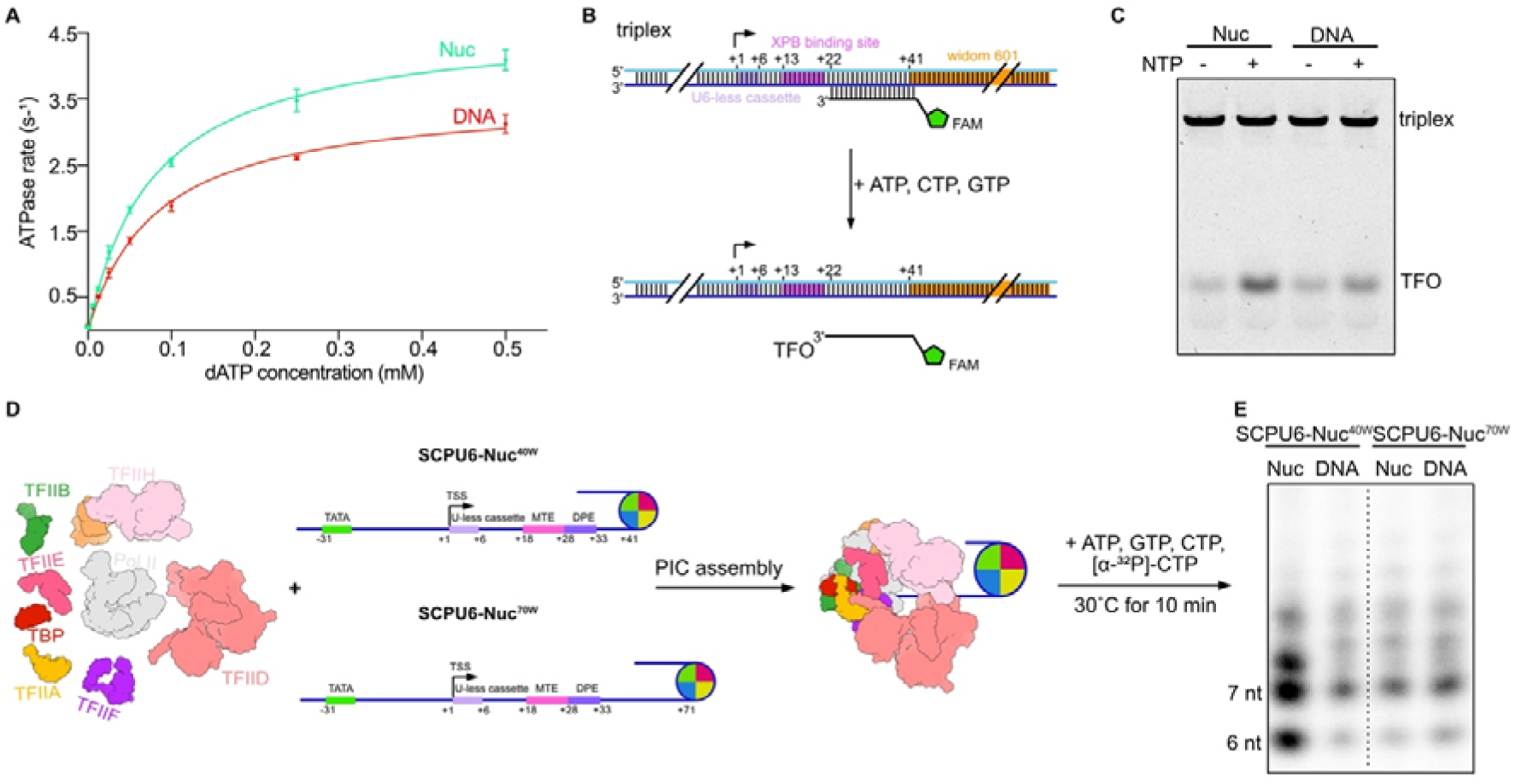
The +1 nucleosome stimulates TFIIH activity and initial transcription. **(A)** NADH-coupled dATPase assay to monitor TFIIH ATPase activity within the PIC in the presence and absence of the +1 nucleosome. The ATPase rates of TFIIH were measured at different concentrations of dATP and fitted with a Michaelis-Menten model (Methods). The experiments were repeated three times, data points are mean ± standard deviation (SD). Nuc, nucleosome. **(B)** Schematic of the triplex-displacement assay. The triplex template contains the U6-less cassette (light purple) and the widom 601 sequence (orange) to position the +1 nucleosome 40 bp from the TSS. The XPB binding region on the triplex in the closed PIC is highlighted in pink. The FAM-labelled TFO is 22 nt long and bound to the DNA duplex from +22 to +43 downstream of the TSS. Upon TFIIH translocation of the DNA template, the duplex-bound TFO is displaced from the triplex. Positions of core XPB binding, U-less cassette and the edge of the +1 nucleosome are indicated relative to the TSS. **(C)** Triplex-displacement assay described in (**B)** to monitor the translocase activity of TFIIH within the PIC, in the presence (Nuc) and absence (DNA) of the +1 nucleosome. The reactions were analysed by native-PAGE and bands corresponding to the triplex and free-TFO are labelled. The experiments were repeated three times. **(D)** Scheme of *in vitro* transcription initiation assay to monitor the influence of the +1 nucleosome on initial transcription with two different nucleosome-TSS distances. Both promoter templates contain a U6-less cassette (light purple) but have the +1 nucleosome positioned 40 and 70 base-pairs downstream of the TSS in SCPU6-Nuc^40W^ and SCPU6-Nuc^70W^, respectively. Positions of core promoter motifs, U-less cassette and the edge of the +1 nucleosome are indicated relative to the TSS. **(E)** Urea-PAGE analysis of the *in vitro* transcription initiation assays using SCPU6-Nuc^40W^ and SCPU6-Nuc^70W^ promoter DNA. Experiments were done with a +1 nucleosome reconstituted (Nuc) and on linear DNA (DNA). Experiments were repeated three times.

In summary, a +1 nucleosome positioned 40 bp downstream of the TSS positively regulates the ATPase and translocase activity of TFIIH as well as initial RNA synthesis.

### The +1 nucleosome facilitates TFIIH eviction during promoter escape

After promoter DNA melting and initial transcription, the ITC is converted to an early EC by upstream bubble rewinding, which coincides with the removal of a subset of GTFs and formation of a stable transcription complex^11–14,16,17^. However, the role of the +1 nucleosome in the ITC-EC transition remains unclear.

To investigate the role of the +1 nucleosome in this transition, we reconstituted the +1 nucleosome on a DNA template containing a U-less cassette that stops transcription after 10 nt in the absence of UTP (Figure 4A), as the transition was shown to occur after the synthesis of 10-14 nt of RNA^16,17,46,47^. We then assembled the PIC, started transcription by adding ATP, CTP and GTP and analysed the reaction mixture by cryo-EM (Figure 4A). From this we obtained a reconstruction of the transcription complex at an overall resolution of 3.9 Å (Figure 4B, Figure S5A-B), where the Pol II active site could be resolved to 3.6 Å. The cryo-EM density of the RNA in the Pol II active site shows that the EC had stalled at 10 nt as expected (Figure 4B, insert), and we therefore call the structure EC10-Nuc^40W^. The overall structure of the EC10-Nuc^40W^ is similar to early EC structures reported before^16,17^ (Figure 4B-C). Whereas the EC10-Nuc^40W^ reconstruction showed density for the +1 nucleosome, density for TFIIH was absent (Figure S5C), contrary to the early EC structures determined on a linear DNA template with a U-less cassette of 14 nt that showed density for TFIIH (EC14)^16^ (Figure 4C). When we superimposed these structures of early ECs, the nucleosome in the EC10-Nuc^40W^ largely overlaps with the position of TFIIH in the EC14^16^ (Figure 4D). This suggests that the +1 nucleosome can facilitate eviction of TFIIH from the transcription complex during promoter escape.

**Figure 4.**
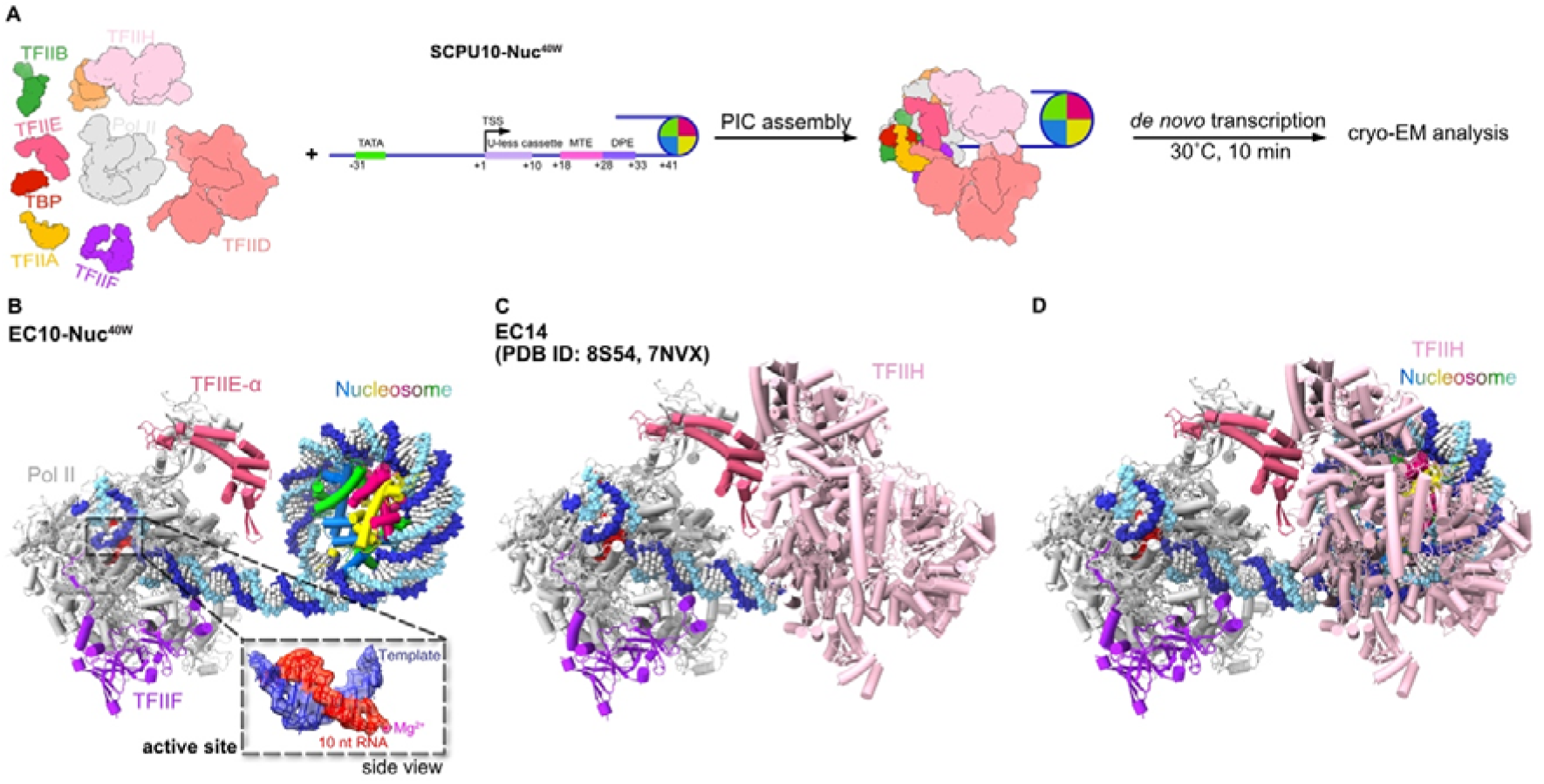
The +1 nucleosome facilitates TFIIH eviction during promoter escape. **(A)** Scheme of the experimental setup for cryo-EM sample preparation of the EC10-Nuc^40W^ structure shown in panel (**B)**, using the SCPU10-Nuc^40W^ promoter DNA. Positions of core promoter motifs, U-less cassette and the edge of the +1 nucleosome are indicated relative to the TSS. **(B)** Structure of the EC10-Nuc^40W^ shown as cartoon in top view, DNA is shown as spheres. The inset depicts the cryo-EM density carved around the Pol II active site showing that Pol II has transcribed 10 nt RNA in the EC10-Nuc^40W^. The active site Mg^2+^ is shown as magenta sphere. **(C)** Structural model for the EC14 in the absence of a +1 nucleosome modelled by fitting PDB ID 8S54^16^ and 7NVX^38^ into the cryo-EM density of EC14 (EMD-19725^16^). **(D)** Superimposition of the EC10-Nuc^40W^ and the the EC14 model showing that TFIIH clashes with the +1 nucleosome.

## DISCUSSION

Previous studies have shown that the +1 nucleosome influences PIC assembly^30,31,34^ and promoter-proximal pausing^50,51^. It is also known how the Pol II elongation complex progresses through a nucleosome^52–57^. However, how transcription initiation and the transition to elongation occur in the context of a nucleosome remains poorly understood both functionally and structurally. With the use of biochemical and structural data, we present a molecular model of how transcription initiates and proceeds in the presence of the +1 nucleosome (Movie S1 and Figure 5). First, after PIC assembly on the promoter, ATP binding to TFIIH alters DNA trajectory and displaces TFIID from the MTE-DPE motif. Second, as initial RNA synthesis occurs, the +1 nucleosome contacts TFIIH and stimulates TFIIH translocase activity and RNA synthesis. Third, further transcription after bubble rewinding moves the +1 nucleosome near the early EC, leading to eviction of TFIIH.

**Figure 5.**
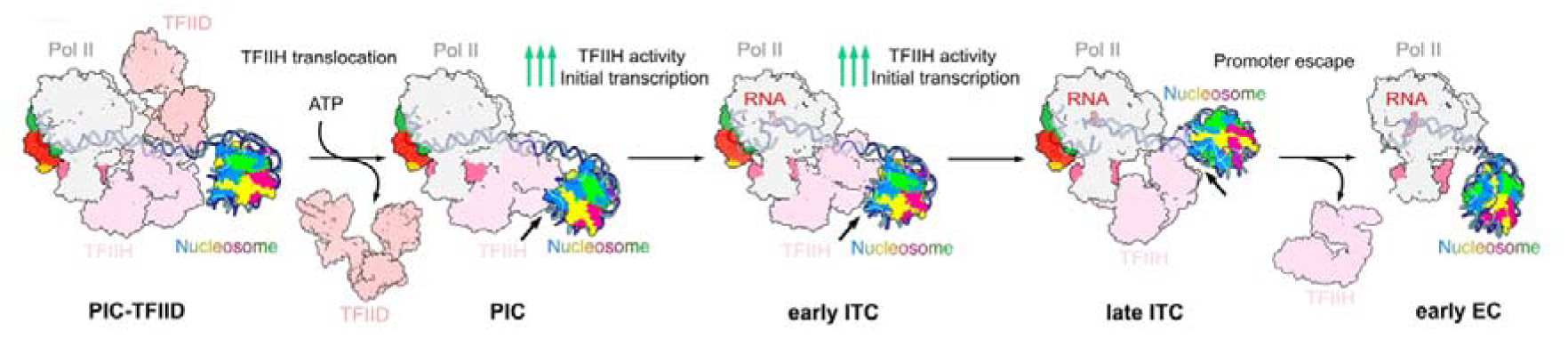
Model of transcription initiation and the transition from initiation to elongation in the presence of the +1 nucleosome. First, in the presence of the +1 nucleosome, TFIID is displaced from the PIC upon the binding of ATP (or ADP-BeF_3_) to TFIIH during promoter melting. Second, the +1 nucleosome interacts with TFIIH (highlighted with an arrow) during promoter opening and initial transcription, stimulating TFIIH activity and thereby initial transcription. Third, upon bubble rewinding, the +1 nucleosome facilitates TFIIH removal from the transcription complex during promoter escape.

Previously determined cryo-EM structures of TFIID-containing PICs assembled on linear promoter DNA show that in the absence of an MTE element, only XPB fully binds promoter DNA downstream the TSS^43^. This suggests that without an MTE motif, affinity of TAF1/7 to the promoter is reduced, allowing XPB to outcompete TFIID from fully binding downstream DNA. However, when the promoter contains an MTE-DPE motif, both XPB and TAF1/7 were observed to bind to the promoter and it is unclear how promoter DNA is handed over from TFIID to TFIIH for transcription initiation. Moreover, in the absence of a +1 nucleosome, TFIID can remain attached to or rebind the promoter region during the initiation-elongation transition^17^. Our structures explain how TFIID is removed from downstream promoter DNA. Binding of ATP (or ADP-BeF_3_) to XPB closes the ATPase lobes on DNA which moves the promoter DNA away from TAF1/7. Additionally, the +1 nucleosome contacts TFIIH and thereby stabilizes the new trajectory of the promoter DNA, which prevents TAF1/7 from rebinding. Retention of TFIID on the transcription complex can slow down transcription^58^, suggesting that when TFIID remains bound it may serve as a roadblock for TFIIH translocation and transcription by Pol II. Thus, destabilization of TFIID by ATP binding to XPB in the presence of a +1 nucleosome, which may eventually lead to TFIID eviction from the PIC, is likely instrumental to facilitate efficient transcription initiation.

We show that the +1 nucleosome enhances TFIIH activity as well as initial transcription, and our structures also suggest the mechanism underlying this stimulation. In the presence of ADP-BeF_3_, DNA binding and translocation by TFIIH lead to a rotation of the +1 nucleosome, which then allows the +1 nucleosome to bind the p52 subunit of TFIIH. p52 has been reported to regulate ATPase activity of XPB^48,49^ and binding of the DNA repair factor XPA to p52 was proposed to positively regulate DNA translocation by XPB^59^. We therefore suggest that binding of the +1 nucleosome to p52 during promoter DNA opening and initial transcription may stabilize the interaction between p52 and XPB, which has been shown to stimulate XPB ATPase activity^48,49^. In addition, XPB lacks processive translocase activity and tends to dissociate from the DNA^8,60^. Binding of the +1 nucleosome to TFIIH might also restrict mobility of the downstream promoter DNA, which may facilitate more efficient DNA translocation by XPB.

Our findings also extend the current model^9–11,15–18,46,47,61–64^ for promoter escape by Pol II. Based on structures of an early EC with TFIIE and TFIIH in the absence of a +1 nucleosome^16,17^, it has been suggested that both TFIIE and TFIIH may remain attached to the complex until the +1 nucleosome clashes with TFIIH^16^. Here, we show that as early transcription proceeds, the +1 nucleosome indeed may displace TFIIH from promoter DNA. Eviction of TFIIH might also destabilize TFIIE^65^ and facilitate displacement by DSIF and NELF^16^. We therefore propose that the +1 nucleosome facilitates promoter escape by controlling TFIIH eviction.

Recent studies have shown that the position of the +1 nucleosome relative to the TSS influences PIC assembly^31,34^. The nucleosome-TFIIH interface described here was previously observed in PICs with an H2A.Z containing and acetylated nucleosome positioned 40-50 bp downstream of the TSS^34^. Histone modifications (such as H3, H4 acetylation and H3K4 trimethylation) and the histone variant H2A.Z on the +1 nucleosome^22,23,66–68,69^ may be bound by PIC components like TFIID^70,71,72^ or Mediator^34,73–76^, potentially inducing transient proximity between the +1 nucleosome and the PIC. Hence, nucleosomes positioned further downstream than 40 bp relative to the TSS may also transiently shift upstream and thereby positively contribute to transcription initiation and the transition to elongation *in vivo*.

### Limitations of the study

Transcription initiation and the transition to elongation are highly dynamic processes, therefore some transient states and flexible complexes might be filtered out during extensive data processing. Additionally, due to the complexity of the *in vitro* transcription initiation system, the work is only performed in the presence of the +1 nucleosome. In the future, it might be interesting to extend this system to the chromatin context.

## ACKNOWLEDGEMENTS

We thank current and past members of the Cramer laboratory, in particular, U. Neef, P. Rus and T. Schulz for maintaining insect cell culture facility and *Sus scrofa* thymus stocks, C. Dienemann and U. Steuerwald for maintenance of the electron microscopy facility, O. Dybkov for maintaining the radioactivity laboratory, K. Lysakovskaia for providing human cDNA for cloning, M. Lidschreiber, K. Zumer and J. Walshe for critical reading of the manuscript, and members of the Cramer lab for helpful discussions.

## AUTHOR CONTRIBUTIONS

Y.Z. and J.A.-G. initiated and planned the project. Y.Z. designed and performed all functional assays. Y.Z. and J.A.-G. reconstituted nucleosomes for cryo-EM analysis. Y.Z. prepared cryo-EM samples PIC-Nuc^40W^_ADP-BeF3_, ITC2-Nuc^40W^, ITC6-Nuc^40W^ and EC10-Nuc^40W^. J.A.-G. prepared cryo-EM sample of PIC-Nuc^40W^. J.A.-G. established the cloning, expression and purification of the recombinant human TFIID from insect cells. Y.Z., J.A.-G. and C.D. collected cryo-EM data. Y.Z. and J.A.-G. processed cryo-EM datasets and built models. F.G. purified general transcription factors. P.S. and J.A.-G designed initial constructs of the recombinant human TFIID and U.N. assisted in their cloning. C.D. and P.C. supervised research. Y.Z., J.A.-G. and C.D interpreted the data. Y.Z., J.A.-G., C.D. and P.C. wrote the manuscript with input from all authors.

## STAR METHODS

### Resource Availability

#### Lead contact

Correspondence and request for materials should be addressed to C.D.

#### Material Availability

Materials are available from Christian Dienemann upon request under a material transfer agreement with the Max Planck Society.

#### Data and Code Availability

The cryo-EM density maps were deposited in the Electron Microscopy Data Bank (EMDB) and coordinates within the Protein Data Bank (PDB) for PIC-Nuc^40W^ (EMD-XXXX, PDB ID XXXX), PIC-Nuc^40W^_ADP-BeF_ (EMD-XXXX, PDB ID XXXX), ITC2-Nuc^40W^ (EMD-XXXX, PDB ID XXXX), ITC6-Nuc^40W^ (EMD-XXXX, PDB ID XXXX)_, and_ EC10-Nuc^40W^ (EMD-XXXX, PDB ID XXXX).

### METHODS DETAILS

#### Protein expression and purification

*S. scrofa* Pol II and human initiation factors (TFIIA, -IIB, -IIE, -IIF, core TFIIH, CAK, CAK-D137R mutant) were expressed and purified as previously described^31,38,59,77,78^ (Figure S1A).

Human TFIID was cloned for expression in insect cells in different 438-series vectors, each representing a different lobe of TFIID. First, we amplified the sequence coding for every subunit from human cDNA by PCR, and combined them by several rounds of ligation-independent cloning (LIC). For initial replication of the constructs, XL1 blue competent cells were employed (Agilent), however, for constructs exceeding 20 kb in size, we replicated them in NEB^®^ Stable Competent E. coli (NEB). The sequences coding for lobe C (TAF1, TAF2 and TAF7), lobe B (TAF4, TAF5, TAF6, TAF8, TAF9, TAF10 and TAF12) and lobe A subunits (TAF3, TAF4, TAF5, TAF6, TAF9, TAF10, TAF11, TAF13 and TBP) were placed in three different vectors. Both TAF2, TAF3 and TAF5 were N-terminally tagged with a 6xHis-maltose binding protein (MBP) tag, followed by a tobacco etch virus (TEV) cleavage site. V1 viruses were generated separately for all three constructs in Sf21 (*Spodoptera frugiperda*).

For its recombinant expression, 600 ml of Hi5 cells (*Trichoplusia Ni*) were co-infected with V1 of lobe A and B, whereas V1 of lobe C was employed alone to infect 3.6 L of Hi5 cells. Cells were harvested separately after 4 days in resuspension buffer (50 mM KOH-HEPES pH 7.5, 300 mM KCl, 10% glycerol, 5 mM MgCl_2_, 5 mM ATP, 1 mM DTT, 0.284 μg/ml leupeptin, 1.37 μg/ml pepstatin A, 0.17 mg/ml PMSF, 0.33 mg/ml benzamidine and 1 tablet of cOmplete™ Protease Inhibitor cocktail (Roche)). Resuspended cell pellets were flash-frozen in liquid nitrogen and stored at −80°C until use. Subsequently, frozen pellets were thawed in a water bath at room temperature, sonicated using a BRANSON digital sonifier for 3 minutes at 30% amplitude in intervals of 0.5 s pulses. The sample was then centrifuged at 87,207 xG for 60 minutes at 4°C to remove the debris, and the resulting supernatant filtered first through a 5 µm filter and then through a 0.8 µm filter. The lysate was loaded on a self-packed XK16/20 column (Cytiva) with 25 ml of amylose resin (New England Biolabs), pre-equilibrated in buffer A1 (50 mM KOH-HEPES pH 7.5, 300 mM KCl, 10% glycerol, 2 mM MgCl_2_, 0.1 mM EDTA, 1 mM DTT, 0.284 μg/ml leupeptin, 1.37 μg/ml pepstatin A, 0.17 mg/ml PMSF and 0.33 mg/ml benzamidine). The colum was washed with buffer A1 and eluted with buffer B1 (50 mM KOH-HEPES pH 7.5, 300 mM KCl, 10% glycerol, 2 mM MgCl_2_, 0.1 mM EDTA, 1 mM DTT, 0.284 μg/ml leupeptin, 1.37 μg/ml pepstatin A, 0.17 mg/ml PMSF, 0.33 mg/ml benzamidine and 100 mM maltose). The elution fractions were pooled and diluted with 0-salt buffer (50 mM KOH-HEPES pH 7.5, 10% glycerol, 2 mM MgCl_2_, 0.1 mM EDTA, 1 mM DTT, 0.284 μg/ml leupeptin, 1.37 μg/ml pepstatin A, 0.17 mg/ml PMSF and 0.33 mg/ml benzamidine) in order to reach a final KCl concentration for the sample of 250 mM KCl. Sample was then incubated with 3-5 mg of TEV protease for 14 hours at 4°C, after which it was injected into a MonoQ 5/50GL column (GE Healthcare), pre-equilibrated with buffer 80% C1 (50 mM KOH-HEPES pH 7.5, 100 mM KCl, 10% glycerol, 2 mM MgCl_2_ and 1 mM TCEP) and 20% buffer C2 (50 mM KOH-HEPES pH 7.5, 1 M KCl, 10% glycerol, 2 mM MgCl_2_ and 1 mM TCEP). This guarantees that the excess of TFIID submodules elute as flow-through and wash fractions. Finally, the column is eluted using a linear gradient from 20% buffer C2 to 100% buffer C2, where fractions containing the stoichiometric TFIID complex were pooled and concentrated using an Amicon 100,000 MWCO Ultra Centrifugal Filter (Merck Millipore). During concentration, the buffer was exchanged with buffer C1, the protein concentrated to 4-5 mg/ml, flash-frozen and stored at −80°C until use (Figure S1A).

#### Nucleosome reconstitution

Super core promoter scaffolds with different U-less cassettes, as well as a 147 bp Widom-601 sequences located at different positions from the TSS, were inserted into pUC119 vectors as previously described^16,30,31,38,79^. A 16 bp extranucleosomal DNA located at the 3’-end of the DNA was included for each DNA template. DNA templates were amplified by large-scale PCR, followed by ion-exchange chromatography and isopropanol precipitation. The purification of *X. laevis* histones, refolding of histone octamers and nucleosome reconstitution were carried out as described^80^. The quality of nucleosome reconstitution was checked on a 1% agarose native gel (0.5x TBE running buffer). Nucleosomes were concentrated and stored at 4°C until use.

#### Cryo-EM sample preparation of transcription intermediates with the +1 nucleosome

Before PIC assembly, the upstream complex TFIID-TFIIA-TFIIB-nucleosome (with a molar ratio 1.4:6.2:3.1:1 for TFIID:TFIIA:TFIIB:nucleosome) was assembled and incubated at room temperature for 30-60 min. The other two subcomplexes, Pol II-TFIIF and TFIIH-TFIIE were assembled in a molar ratio of 1:5 and 1:1, respectively, and incubated at room temperature for 5-10 min. Afterwards, the three subcomplexes were mixed together such that TFIIH and TFIIE would be in a 1.5x molar excess over Pol II, and incubated at 30°C for 60 min in buffer containing 20 mM K-HEPES pH 7.5, 109 mM KCl, 3% glycerol, 6 mM MgCl_2_, 0.5 mM DTT.

PIC-Nuc^40W^ and PIC-Nuc^40W^_ADP-BeF3_ were prepared as described above with the exception that 0.5 mM ADP-BeF_3_ was added to PIC-Nuc^40W^_ADP-BeF3_ during PIC assembly and incubated at 30°C for 30 min.

For the ITC2-Nuc^40W^ sample, transcription was initiated with 0.5 mM ATP and 0.5 mM CTP and incubated for 10 min at 30°C. For the ITC6-Nuc^40W^ and the EC10-Nuc^40W^ samples, transcription was initiated by adding 0.5 mM ATP, 0.5 mM GTP and 0.5 mM CTP and allowed to proceed for 10 min at 30°C.

Afterwards, the complexes were loaded onto a 15-40% (w/v) sucrose gradient while mildly crosslinked by GraFix^81^ at 32,000 rpm (SW60) for 16 hr at 4°C, by supplementing the 40% sucrose solution with 0.05-0.1% glutaraldehyde. For the ITC2-Nuc^40W^ sample, Grafix was perfomed at 55,000 rpm (TLS55) for 3 hr at 4°C. After ultracentrifugation, the gradients were fractionated manually from the top and immediately quenched with 10 mM lysine and 40 mM aspartate for 10 min at 4°C. Fractions of interest were pooled and dialysed against dialysis buffer containing 20 mM K-HEPES pH 7.5, 90 mM KCl, 1% glycerol (v/v) and 1mM DTT at 4°C for 7-8 h to remove sucrose.

For grid preparation, a thin layer of homemade continuous carbon film (∼2.5 nm) was floated onto the dialysed samples and incubated at 4°C for 15 min to absorb the particles onto the film. Afterwards, the film was picked up by a holey-carbon grid (Quantifoil R3.5/1, copper, mesh 200) and washed with 3.5 uL of dialysis buffer. The grids were blotted for 1.5 s with blot force of 5 and plunge-frozen in liquid ethane with a Vitrobot Mark IV (FEI) operated at 4°C and 100% humidity.

#### Cryo-EM data collection and data analysis

Cryo-EM data was collected with an FEI Titan Krios transmission electron microscope equipped with a K3 summit direct electron detector (Gatan) and a GIF quantum energy filter (Gatan). The microscope was operated at 300 keV and with a slit width of 20 eV. Automated data acquisition was carried out with SerialEM^82^ at a nominal magnification of 81,000x (1.05 Å per pixel) with a total dose of approximately 40 e/Å^2^, fractionated over 40 frames. Images were collected with a defocus range of 0.7 µm to 1.7 µm.

Warp^83^ was used for on-the-fly motion correction, contrast-transfer function estimation and particle picking, with the exception of PIC-Nuc^40W^, for which it was done using cryoSPARC^84^. The datasets were first cleaned by consecutive rounds of 2D and 3D classifications in cryoSPARC^84^ to remove ice contamination, falsely picked and aggregated particles. Particles with good densities for Pol II were then imported into Relion3.1 or Relion5.0^85,86^ for further processing, as detailed in Extended Data Fig. 1, 3 and 5.

For PIC-Nuc^40W^, after initial cleaning in cryoSPARC, particles were further classified by 2D classification in RELION (Figure S1B). Subsequently, focused classifications with signal subtractions were carried out for TFIIH-TFIID, TFIIH and the nucleosome alone. Particles with good density for the nucleosome were reverted and a focused refinement was performed to improve the resolution for cPIC. These particles were also intersected with the improved map of TFIIH-TFIID to generate a consensus map with better density for this part of the structure.

For PIC-Nuc^40W^_ADP-BeF_, the cleaned-up particles were first subtracted with a mask around TFIIH and nucleosome and subjected to one round of global 3D classification (Figure S1C). The subtracted particles with good densities for TFIIH were pooled and reverted to original particles. The reverted particles were then CTF-refined and polished. The resolutions of the nucleosome, TFIIH and XPB were improved by signal subtraction followed by 2D or 3D classifications and focused refinements. Particles with good densities for the nucleosome were intersected with particles with good densities for TFIIH, generating a reconstruction for PIC-Nuc^40W^_ADP-BeF_.

For ITC2-Nuc^40W^ dataset, the clean-up particles were subjected to one round of focused 3D classification with a mask around cPIC to separate closed PIC and ITC complexes (Figure S3A). To improve the resolutions of the nucleosome, TFIIH and TFIIH-nucleosome, the ITC particles signal subtracted and subjected to rounds of classifications and refinements. To improve the resolution of cITC, the ITC particles were CTF-refined, polished, focused 3D classified and focused 3D refined. To obtain a reconstruction of the ITC2-Nuc^40W^, subtracted particles with good densities for TFIIH-nucleosome were reverted back to the original particles and 3D refined.

For ITC6-Nuc^40W^, the cleaned-up particles were separated in CC and ITC-like complexes by performing 3D classification without image alignment with a mask on cPIC (Figure S3B). First, these particles were separately CTF-refined and polished. For CC complexes, a series of focused masked classifications with signal subtraction were performed to improve the resolution of the different part of the complexes. For ITC-like complexes, signal subtraction with a mask on cITC was carried out to perform 3D classifications without image alignment to separate ITC with different lengths of RNA. Through this, we obtained an EC of 9 nt, an ITC of 8 nt and an ITC of 6 nt. For these same ITC-like complexes, a series of global 3D classifications with alignment were performed to filter out those states lacking TFIIH (cITC-Nuc). For the particles containing TFIIH, signal subtraction procedures followed by masked classifications were performed to improve the resolution of TFIIH and the nucleosome. This set of particles was then reverted to obtain the consensus map of the ITC6-Nuc^40W^. Additionally, focused refinement of this consensus around cITC showed clear density for a RNA chain of 6 nt at the active site of Pol II.

For EC10-Nuc^40W^, the cleaned-up particles were CTF-refined and polished (Figure S5C). The particles were then subtracted with a mask around the nucleosome and subjected to multiple rounds of 2D and 3D classifications in cryoSPARC^84^ to improve the resolution of the nucleosome. The subtracted particles with good densities for nucleosome were imported back to Relion5.0, reverted back to the original particles and 3D refined to obtain a reconstruction of the EC10-Nuc^40W^. Focused 3D refinement was performed to improve the resolution of core Pol II.

#### Model building

To facilitate model building, the cryo-EM maps were filtered based on their local resolutions in Relion3.1 or Relion5.0^85,86^ and auto-sharpened in Phenix^87,88^.

For modelling of PIC-Nuc^40W^, PIC-Nuc^40W^_ADP-BeF_, ITC2-Nuc^40W^, and ITC6-Nuc^40W^, previously published models (PDB ID: 7NW0^38^, 8S51^16^, 7NVX^38^, 8BZ1^31^, 8BVW^31^, 8GXS^34^) were used as starting models. For modelling of the EC10-Nuc^40W^, previously published models (PDB ID: 8S54^16^ and 9J0P^51^) were used as starting models. They were docked into focused refinement maps in Chimera^89^, followed by manual adjustments and connection of the linker nucleosomal DNA in Coot^90^. The resulting model was then subjected to consecutive rounds of real-space refinements in Phenix^87,88,91^ and rebuilding with Coot^90^ and ISOLDE^92^. The final models were subjected to comprehensive validation (cryo-EM) in Phenix^87,88^ and showed good stereochemistry.

#### *In vitro* transcription initiation assay

Transcription initiation assays were performed as previously described^16,38^, with modifications. Promoter templates with and without nucleosome were prepared as described above. In brief, for each 15 uL reaction, 2.4 pmol Pol II, 12.2 pmol TFIIF, 2.6 pmol DNA (or nucleosome), 3.7 pmol TFIID, 16 pmol TFIIA, 8 pmol TFIIB, 3.7 pmol TFIIE and 3.7 pmol TFIIH were used. PIC assembly was performed in the same way as cryo-EM sample preparation. Transcription was initiated by adding an NTP mix containing 0.5 mM ATP, 0.5 mM CTP, 0.5 mM GTP and 2 μCi [α-^32^P]-CTP and allowed to proceed for 10 min at 30°C, in a reaction buffer containing 20 mM K-HEPES pH 7.5, 109 mM KCl, 3% glycerol, 6 mM MgCl_2_, 0.5 mM DTT. Reactions were stopped at 37°C for 30 min by the addition of 42 uL stop buffer, containing a final buffer condition of 10 mM K-HEPES pH 7.5, 50 mM EDTA, 1 mg/mL proteinase K, 0.3 M sodium acetate pH 5.5 and 0.8 mg/mL GlycoBlue. The RNAs were isopropanol precipitated and loaded onto an urea-gel (7 M urea, 1x TBE, 20% acrylamide:bis-acrylamide 19:1).

#### Triplex-displacement assay

Triplex-displacement assays were performed as described, with modifications^8,93^. A 22-bp TFO binding site was inserted into pUC119-SCPU6Nuc^40W^ vector. The DNA was amplified by large-scale PCR and purified, as described above. For TFO annealing, 1000 pmol FAM-TFO and 1000 pmol DNA were mixed in annealing buffer (40 mM MES pH 5.5 and 10 mM MgCl_2_). The mixture was heated at 57°C for 15 min and then slowly cooled down to room temperature. The annealed triplex DNA was stored at −20°C until use. For nucleosome reconstitution, equimolar amount of triplex DNA and purified *X.laevis* histone octamers were mixed in reaction buffer containing 1.2 M NaCl, 40 mM Na-cacodylate pH 6.0 and 5 mM MgCl_2_ and incubated at 4°C for 30 min. Afterwards, stepwise dialysis was performed against buffer 0.85 CAM (850 mM NaCl, 20 mM Na-cacodylate pH 6.0, 5 mM MgCl_2_), 0.65 CAM (650 mM NaCl, 20 mM Na-cacodylate pH 6.0, 5 mM MgCl_2_), 0.5 CAM (500 mM NaCl, 20 mM Na-cacodylate pH 6.0, 5 mM MgCl_2_) and 0.1 CAM (100 mM NaCl, 20 mM Na-cacodylate pH 6.0, 5 mM MgCl_2_) at 4°C for 2 hr each. Finally, the nucleosomes were dialysed against buffer containing 20 mM K-HEPES pH 7.5, 50 mM NaCl and 5 mM MgCl_2_ at 4°C overnight. The reconstituted nucleosomes were stored at 4°C until use.

A typical 4.5-uL assay was performed with 1.1 pmol Pol II, 5.8 pmol TFIIF, 1.2 pmol DNA/nucleosome, 7.6 pmol TFIIA, 1.7 pmol TBP, 3.8 pmol TFIIB, 1.7 pmol TFIIE and 1.7 pmol TFIIH in final buffer containing 20 mM K-HEPES pH 7.5, 109 mM NaCl, 3% glycerol, 16 mM MgCl_2_ and 0.5 mM DTT. PIC assembly was carried out as previously described for cryo-EM analysis, except that the complex was incubated at 4°C for 2 hr. Reactions were initiated by a mixture of 0.5 mM ATP, 0.5 mM CTP, 0.5 mM GTP and 2 µM unlabelled TFO and incubated at 30°C for 10 min. Reactions were then quenched with stop buffer containing 0.6% SDS and 2 uL proteinase K (NEB, 20 mg/mL) and loaded onto a NuPAGE 3-8% Tris Acetate protein gel (Thermo Scientific). The gels were scanned with Typhoon FLA 9500 (GE Healthcare Life Sciences).

#### dATPase assay

The dATPase assay is based on the coupling of ATP regeneration to NADH oxidation by two fast enzymatic reactions. A typical 30-uL reaction contains 1.8 pmol Pol II, 9 pmol TFIIF, 1.9 pmol DNA/nucleosome, 12 pmol TFIIA, 2.7 pmol TBP, 6 pmol TFIIB, 1.5 pmol TFIIE, 1.5 pmol CAK (kinase dead mutant CDK7-D137R) and 1.5 pmol core TFIIH in assembly buffer (20 mM K-HEPES pH 7.5, 109 mM KCl, 3% glycerol, 6 mM MgCl_2_, 0.5 mM DTT). PIC assembly was carried out as previously described for cryo-EM analysis, and additionally the CDK7 inhibitor SY-5906 was added to a final concentration of 10 µM and the mixture was further incubated at 30°C for 10 min. A mixture of 2 mM phosphoenolpyruvate (PEP), 0.25 mM NADH and excess pyruvate kinase/lactate dehydrogenase enzyme mix was added and the reaction was incubated at 30°C for 10 min. Reactions were initiated by the addition of dATP at different concentrations (0 µM, 6.25 µM, 12.5 µM, 25 µM, 50 µM, 100 µM, 250 µM and 500 µM).

#### Quantification and statistical analysis

For dATPase assays, the decrease in absorption at 340 nm was measured with the Infinite M1000Pro reader (Tecan). Resulting curved were fitted to a linear model with GraphPad Prism (version 10) and ATPase rates were calculated and plotted against dATP concentrations and fitted with a Michaelis-Menten model. The experiments were performed three times. *In vitro* initiation assays and triplex displacement experiment were performed three times.

## SUPPLEMENTARY FIGURE LEGENDS

**Figure S1. Cryo-EM structure determination of PIC-Nuc^40W^ and PIC-Nuc^40W^_ADP-BeF_.**

**(A)** SDS-PAGE analysis of *Sus scrofa* Pol II and human general transcription factors. The gels were stained by Coomassie blue.

**(B)** Cryo-EM data processing scheme of the PIC-Nuc^40W^ (without ADP-BeF_3_) sample. Reconstructions used for model building are highlighted with a box. The particles number and average resolution of each reconstruction are reported next to the densities.

**(C)** Cryo-EM data processing scheme of the PIC-Nuc^40W^_ADP-BeF_ (with ADP-BeF_3_) sample. Reconstructions used for model building are highlighted with a box. The particles number and average resolution of each reconstruction are reported next to the densities.

**Figure S2. Quality of cryo-EM reconstructions of PIC-Nuc^40W^ and PIC-Nuc^40W^**

**(A and B)** All maps are filtered according to local resolution, with their angular distribution indicated. Fourier shell correlation (FSC) of all the maps is shown. Average resolutions reported in **Figure S1B-C** are estimated at the FSC 0.143 cut-off criterion (dashed line).

**(C)** Focused refinement map and model of XPB bound with ADP-BeF_3_.

**(D)** Global refinement maps of the PIC-Nuc^40W^.

**(E)** Global refinement maps of the PIC-Nuc^40W^_ADP-BeF_.

**(F)** The interaction between the p52 subunit of TFIIH and the nucleosome in the PIC-Nuc^40W^_ADP-BeF_. p52 interacts with H3 and H4.

**Figure S3. Cryo-EM structure determination of ITC2-Nuc^40W^ and ITC6-Nuc^40W^**

**(A)** Cryo-EM data processing scheme of the ITC2-Nuc^40W^ sample. Reconstructions used for model building are highlighted with a box. The particles number and average resolution of each reconstruction are reported next to the densities.

**(B)** Cryo-EM data processing scheme of the ITC6-Nuc^40W^ sample. Reconstructions used for model building are highlighted with a box. The particles number and average resolution of each reconstruction are reported next to the densities.

**Figure S4. Quality of cryo-EM reconstructions of ITC2-Nuc^40W^ and ITC6-Nuc^40W^**

**(A and B)** All maps are filtered according to local resolution, with angular distribution indicated. Fourier shell correlation (FSC) of all the maps is shown. Average resolutions reported in **Figure S3 A-B** are estimated at the FSC 0.143 cut-off criterion (dashed line).

**(C)** Global refinement maps of the ITC2-Nuc^40W^ (left) and the ITC6-Nuc^40W^ (right). Colour legends are depicted below the figure.

**(D)** Superimposition of the ITC2-Nuc^40W^ and the PIC-Nuc^40W^_ADP-BeF_ aligning on Pol II RPB1 subunit. Colour legends are depicted below the figure. All components follow the colour scheme used throughout the manuscript, except for TFIIH and the nucleosome.

**(E)** Superimposition of the ITC2-Nuc^40W^ and the ITC6-Nuc^40W^ aligning on Pol II RPB1 subunit. Colour legends are depicted below the figure. All components follow the colour scheme used throughout the manuscript, except for TFIIH and the nucleosome. Pol II and the GTFs are shown in transparent for clarity.

**(F)** The interaction between p52-XPB subunits interface of TFIIH and the nucleosomal DNA in the ITC6-Nuc^40W^. The interface is indicated with an arrow. p52 is shown in purple and XPB is shown in dark pink.

**(G)** Initial and last frame of the multibody refinement performed on the ITC6-Nuc^40W^ for the 3 components used. Eigenvalues for the 3 motion components are shown.

**Figure S5. Analysis of the role of the +1 nucleosome in early elongation.**

**(A)** Cryo-EM data processing scheme for EC10-Nuc^40W^ sample. The reconstructions used for model building are highlighted with a box. The corresponding particle numbers and overall resolution for each reconstruction are indicated.

**(B)** Quality of cryo-EM reconstructions of the EC10-Nuc^40W^. All maps are filtered according to local resolution, with their angular distribution indicated. Fourier shell correlation (FSC) of all the maps is shown. Average resolutions reported are estimated at the FSC 0.143 cut-off criterion (dashed line).

**(C)** Global map of the EC10-Nuc^40W^.

**Table S1.**
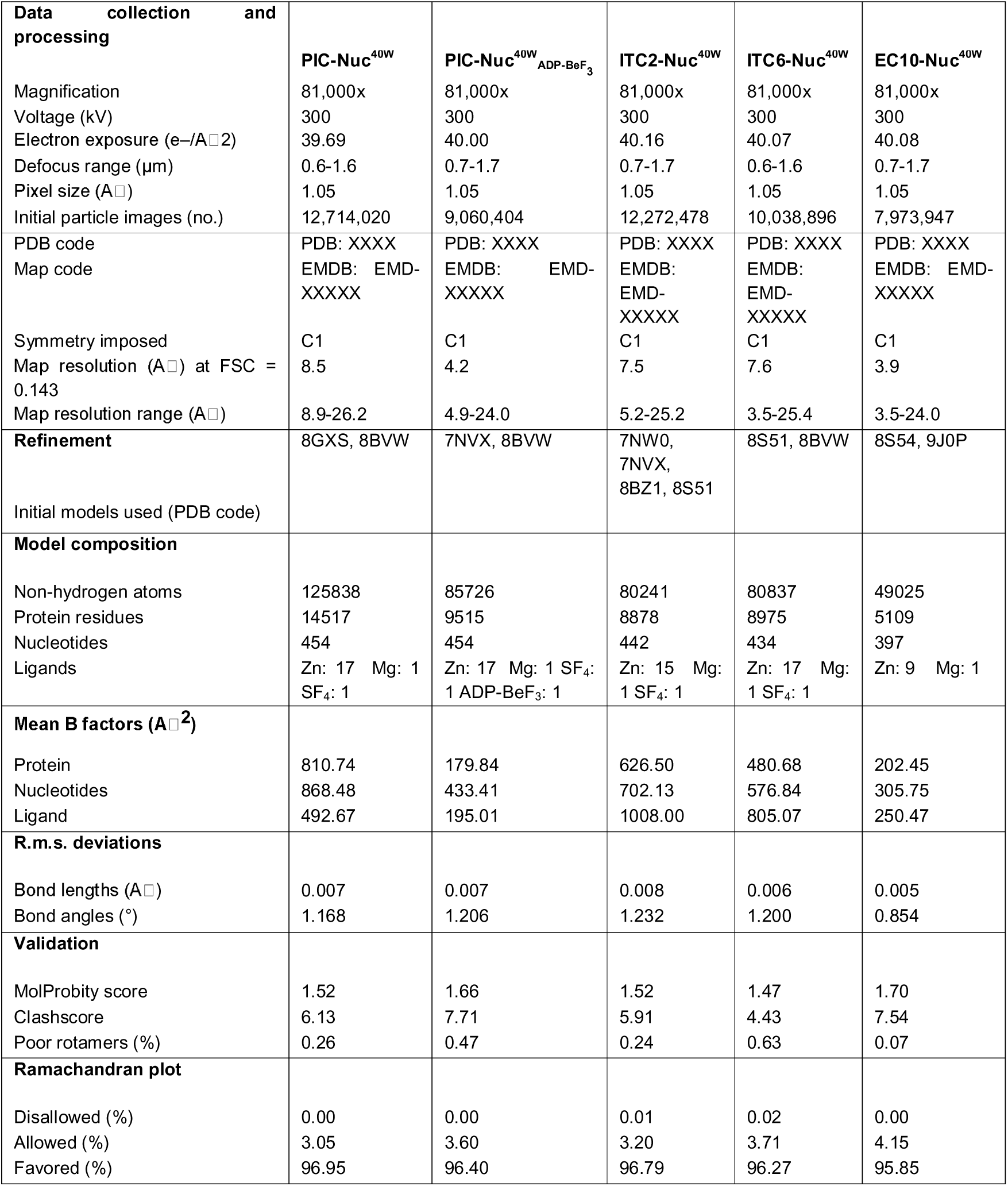
Cryo-EM data acquisition, processing and refinement statistics.

## SUPPLEMENTARY DATA

**Movie S1. The +1 nucleosome functions in Pol II transcription initiation and the transition to elongation.**

This video shows the model of transcription initiation and the transition to elongation in the presence of the +1 nucleosome. First, after PIC assembly, the +1 nucleosome facilitates the eviction of TFIID from the PIC upon binding of ATP to TFIIH. Second, during initial RNA synthesis, the +1 nucleosome contact TFIIH, stimulates TFIIH translocase activity and initial RNA synthesis. Finally, after DNA bubble rewinding, the +1 nucleosome facilitates TFIIH removal from the early elongation complex for promoter escape.

## KEY RESOURCES TABLE

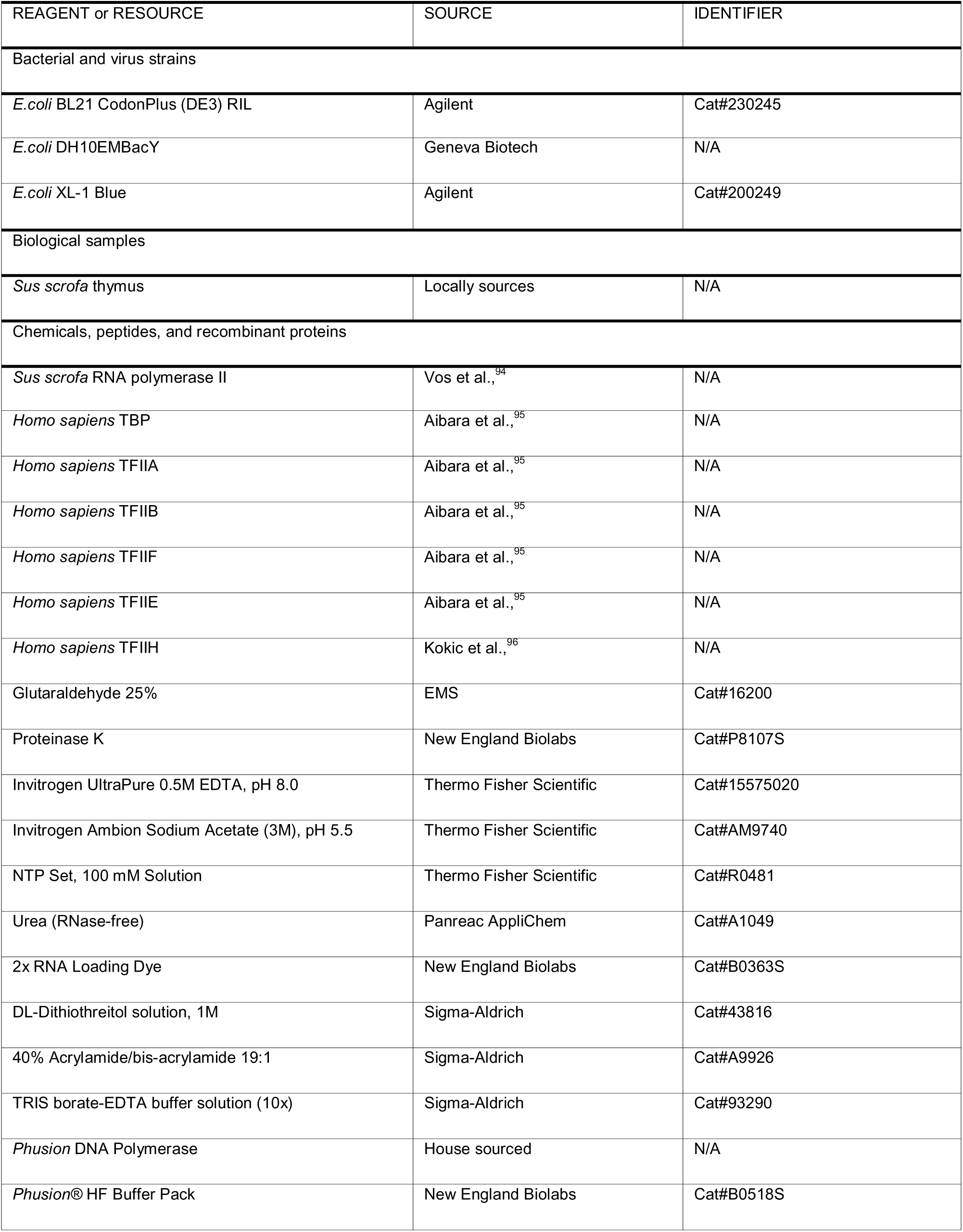
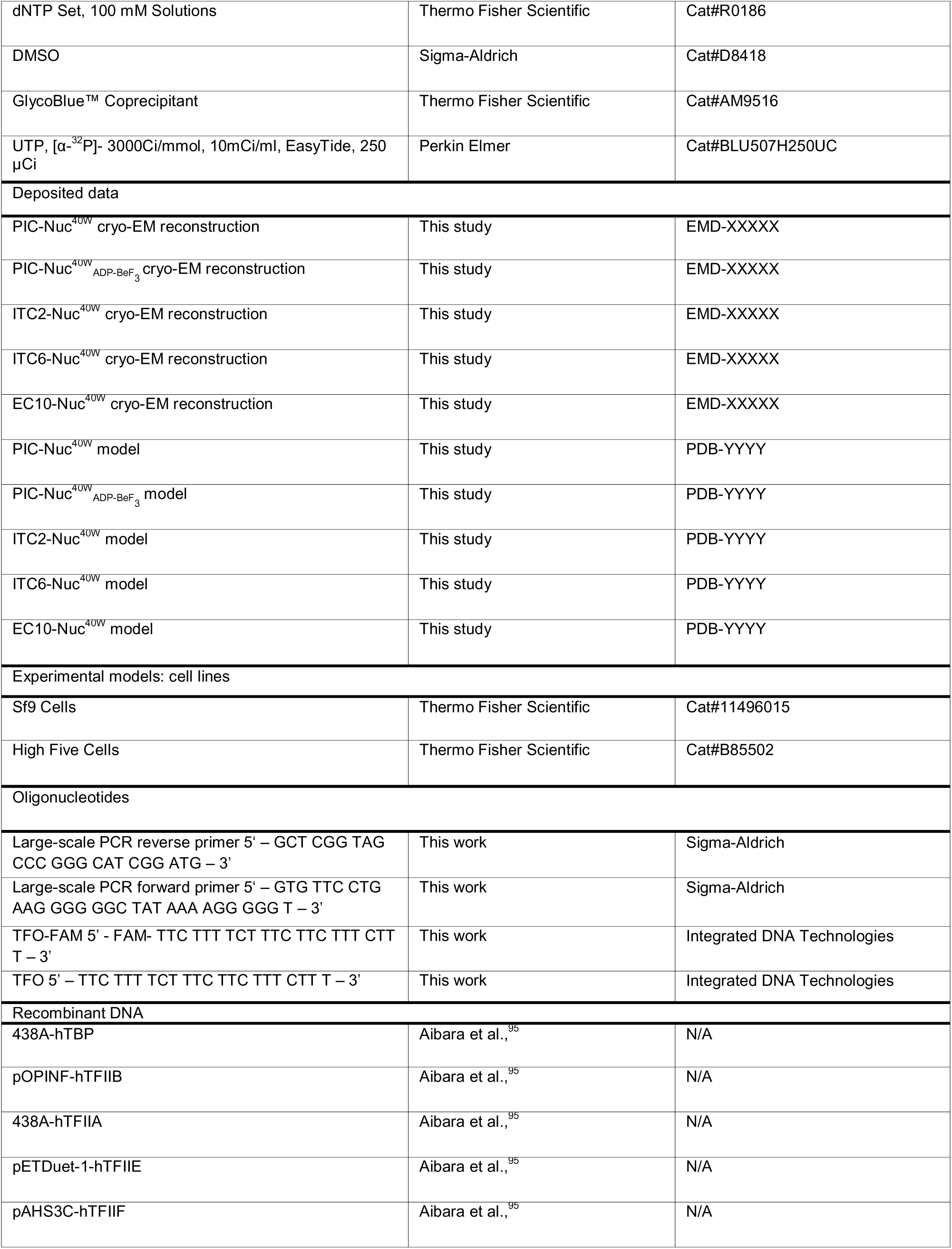
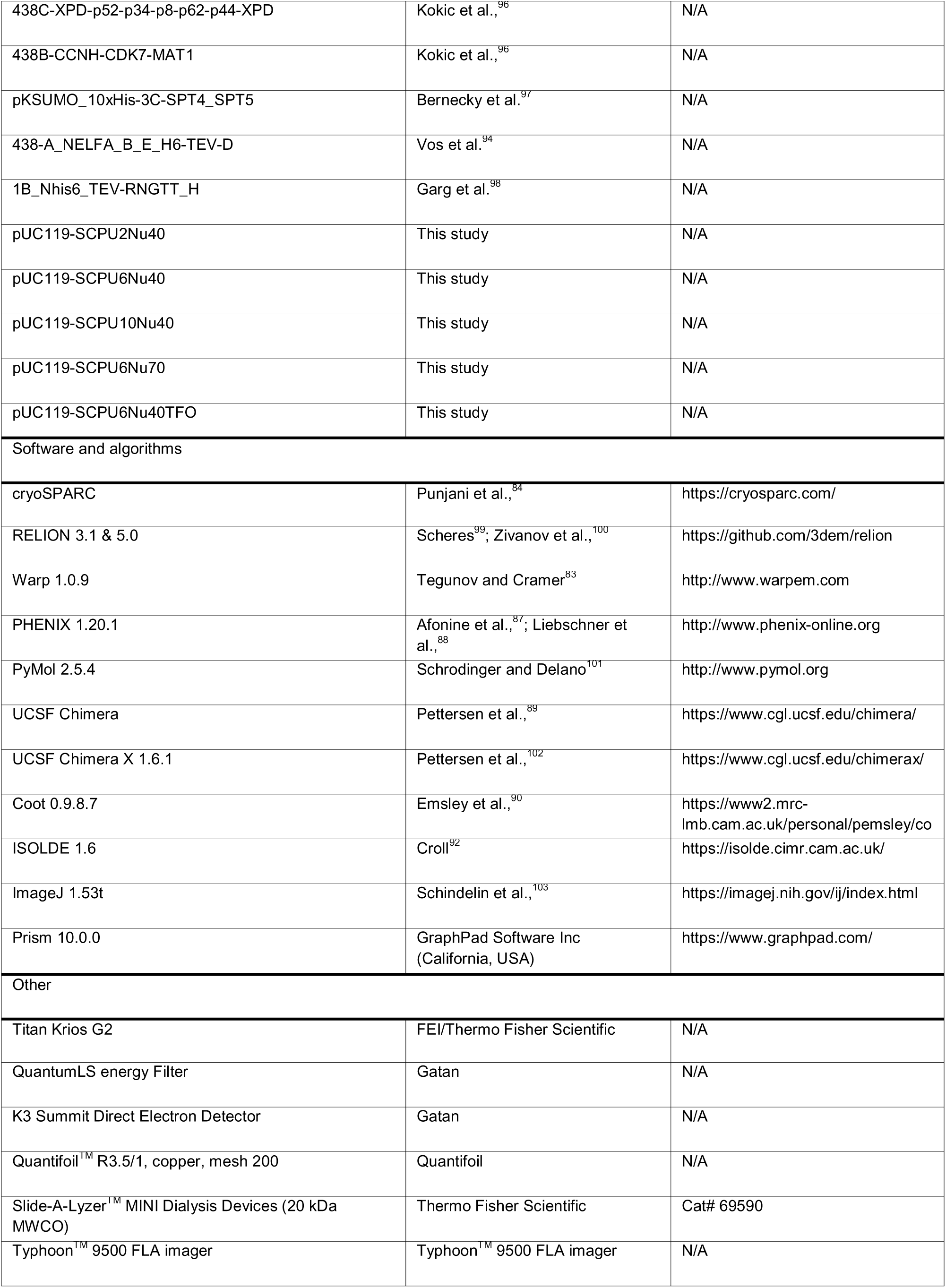

