## Supplementary Figures for "The +1 nucleosome functions in Pol II transcription initiation and the transition to elongation"

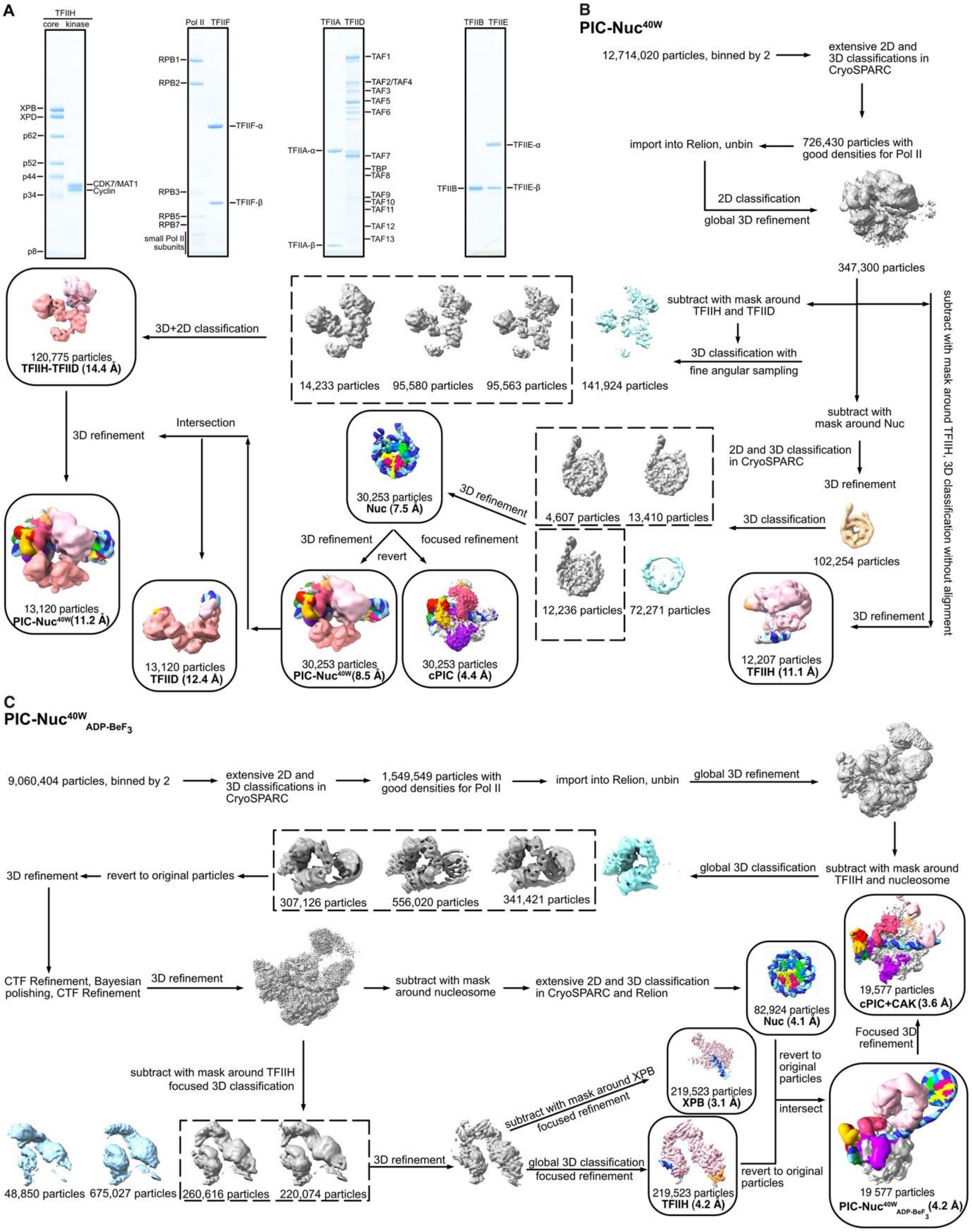


**Figure S1. Cryo-EM structure determination of PIC-Nuc$^{40W}$ and PIC-Nuc$^{40W}_{ADP\text{-}BeF_3}$, related to Figure 1**

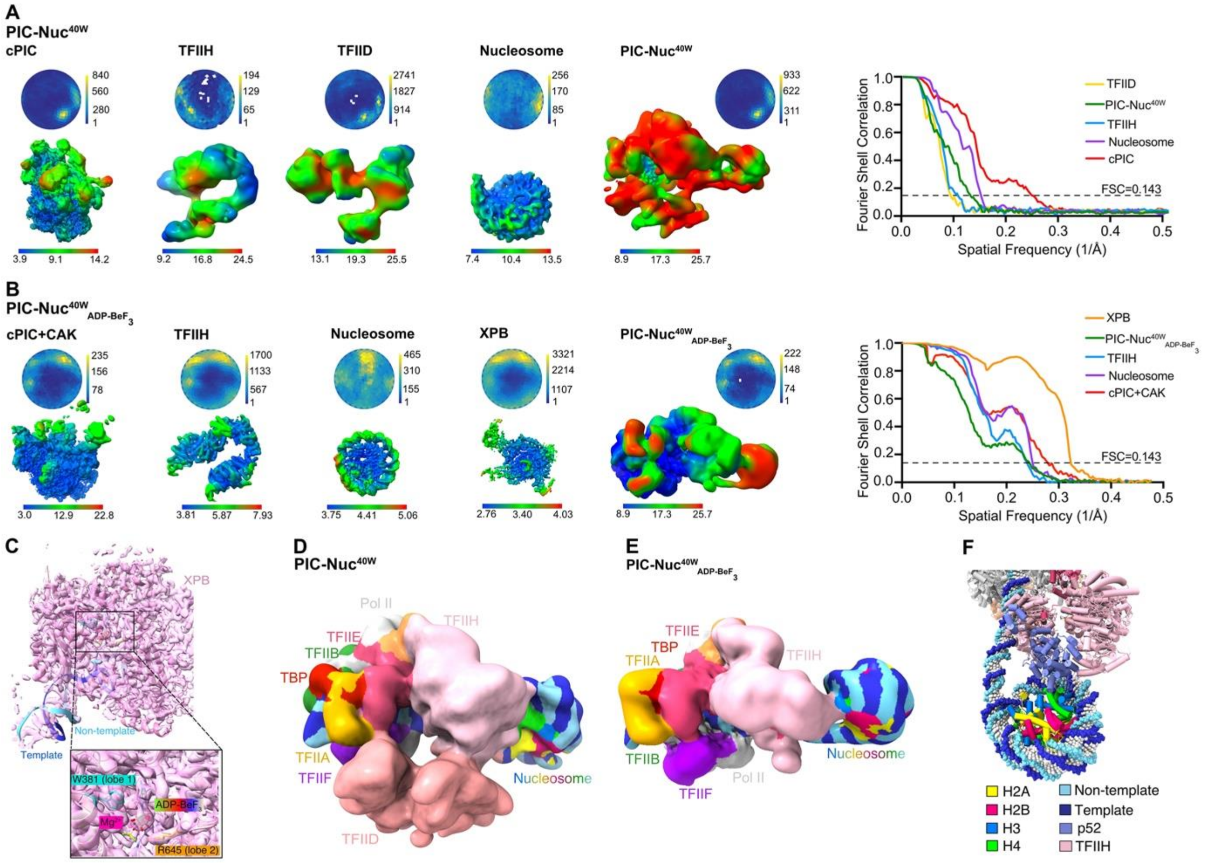

**Figure S2. Quality of cryo-EM reconstructions of PIC-Nuc$^{40W}$ and PIC-Nuc$^{40W}$$_{ADP\text{-}BeF_3}$, related to Figure 1.**

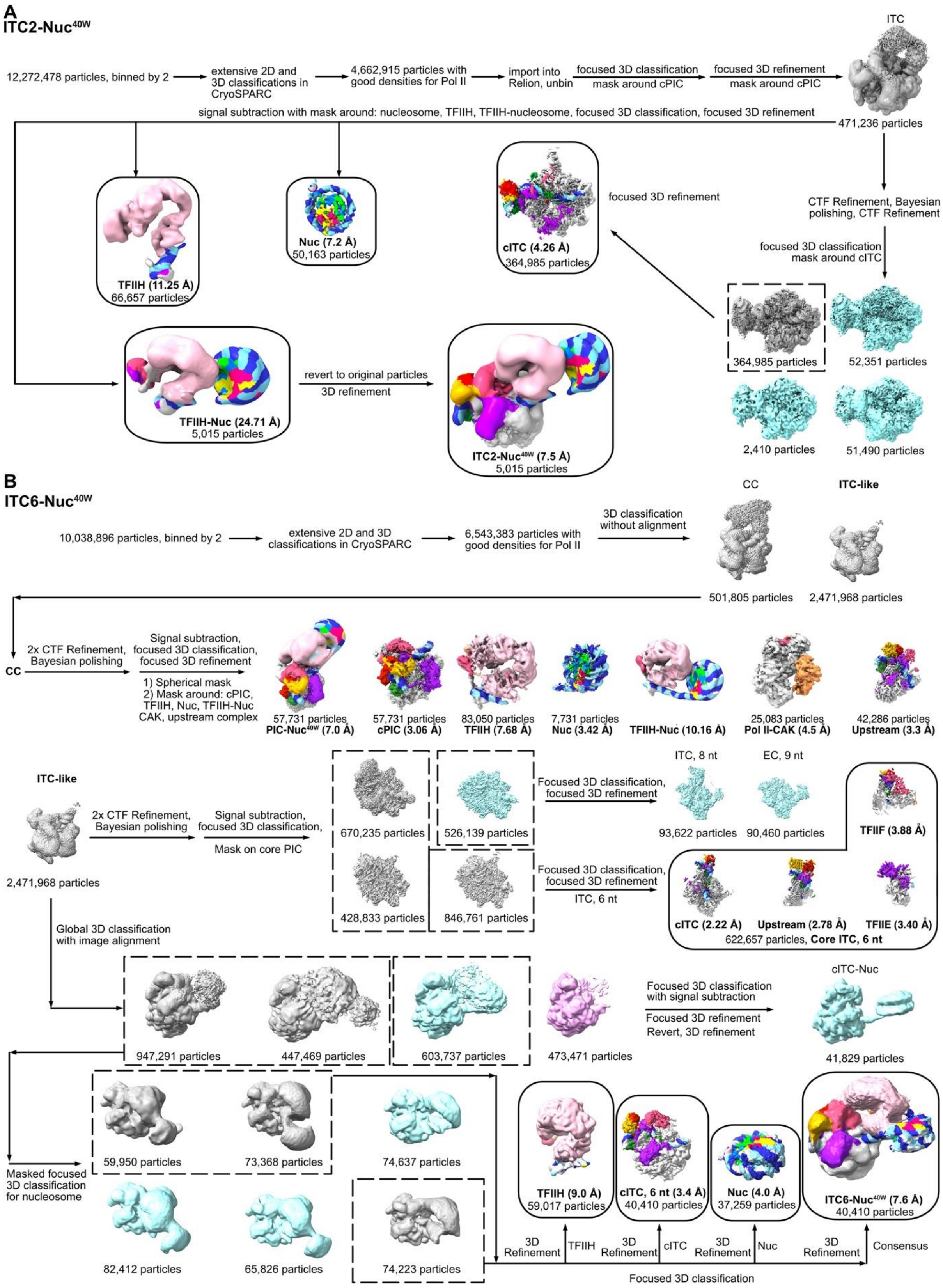


**Figure S3. Cryo-EM structure determination of ITC2-Nuc$^{40W}$ and ITC6-Nuc$^{40W}$, related to Figure 2.**

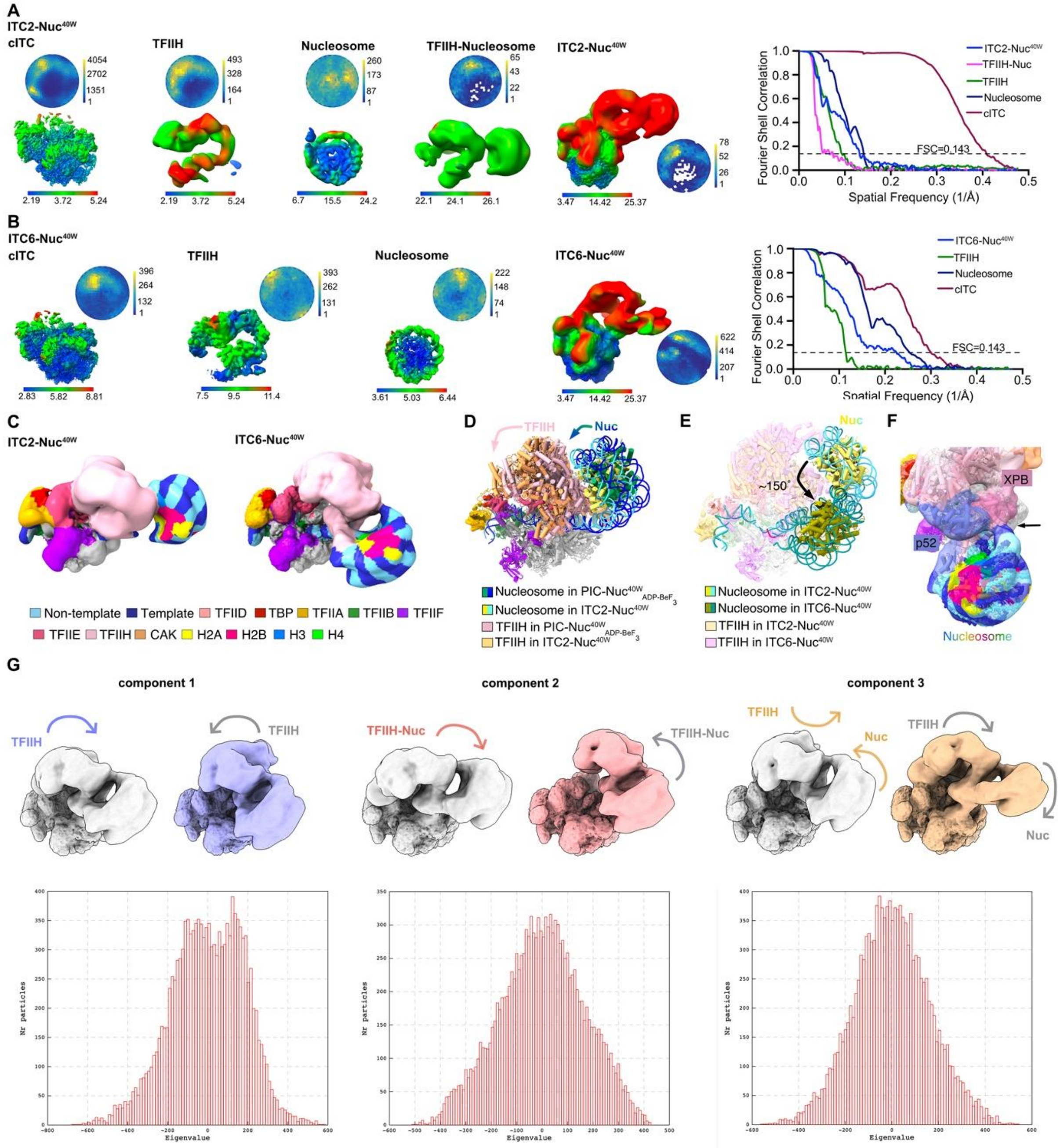


**Figure S4. Quality of cryo-EM reconstructions of ITC2-Nuc$^{40W}$ and ITC6-Nuc$^{40W}$, related to Figure 2.**

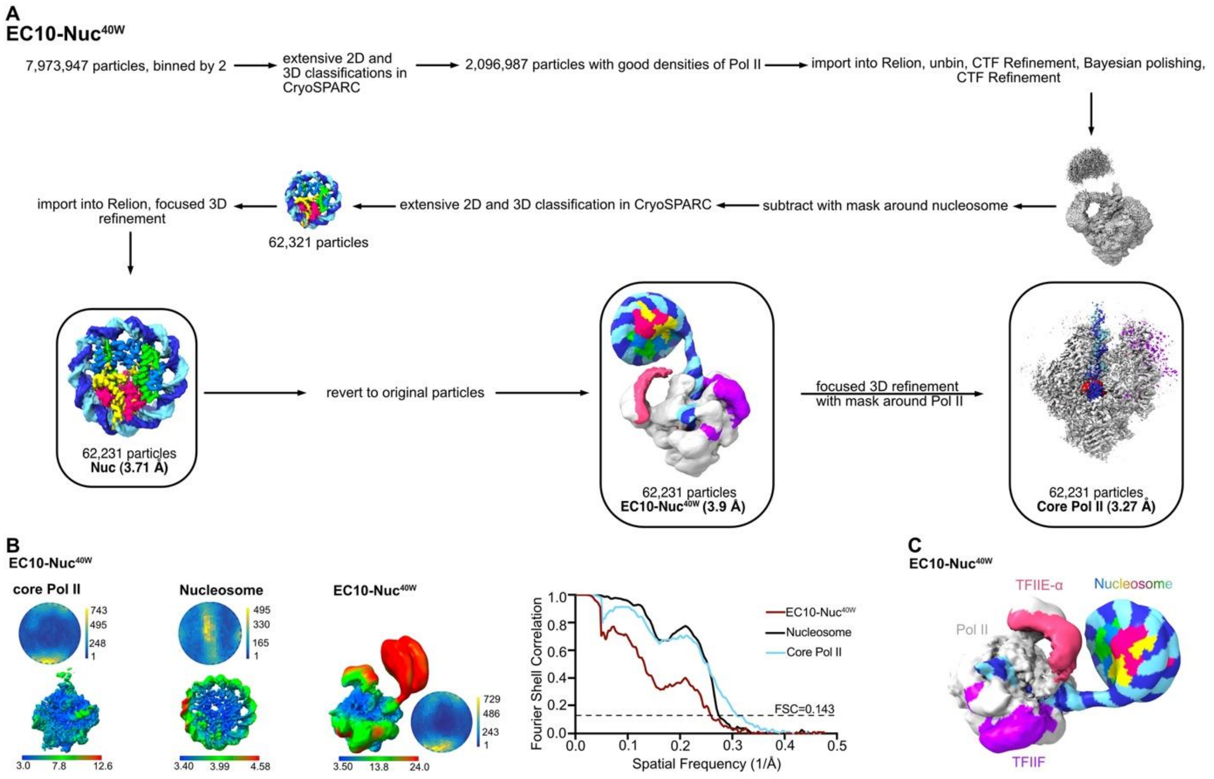


**Figure S5. Analysis of the role of the +1 nucleosome in early elongation, related to Figure 4.**
